# Viral diversity and zoonotic risk in endangered species

**DOI:** 10.1101/2022.06.27.497730

**Authors:** Kayla Nikc, Gregory F. Albery, Daniel J. Becker, Evan A. Eskew, Anna C. Fagre, Sadie J. Ryan, Colin J. Carlson

## Abstract

A growing body of evidence links zoonotic disease risk, including pandemic threats, to biodiversity loss and other upstream anthropogenic impacts on ecosystem health. However, there is little current research assessing viral diversity in endangered species. Here, combining IUCN Red List data on 5,876 mammal species with data on host-virus associations for a subset of 1,273 extant species, we examine the relationship between endangered species status and viral diversity, including the subset of viruses that can infect humans (zoonotic viruses). We show that fewer total viruses and fewer zoonotic viruses are known to infect more threatened species. After correcting for sampling effort, zoonotic virus diversity is mostly independent of threat status, but endangered species—despite a higher apparent research effort—have a significantly lower diversity of viruses, a property that is not explained by collinearity with host phylogeography or life history variation. Although this pattern could be generated by real biological processes, we suspect instead that endangered species may be subject to additional sampling biases not captured by the total volume of scientific literature (e.g., lower rates of invasive sampling may decrease viral discovery). Overall, our findings suggest that endangered species are no more or less likely to host viruses that pose a threat to humans, but future zoonotic threats might remain undiscovered in these species. This may be concerning, given that drivers of endangered species’ vulnerability such as habitat disturbance, wildlife trade, or climate vulnerability may increase virus prevalence in reservoirs and risk of spillover into humans.

## Introduction

The devastating effects of the COVID-19 pandemic are a reminder of the importance of pathogen surveillance for managing and preventing outbreaks of zoonotic disease. Over the last two decades, scientists across a wide variety of fields have increasingly collaborated under the banner of a “One Health” approach, which aims to foster improved environmental, wildlife, and human health as interlinked and synergistic outcomes. Conservation biologists have specifically underscored an emerging body of evidence concerning environmental degradation, loss of biodiversity, and pandemic risk as motivation to enact stronger conservation policies (Evans et al. 2020; Johnson et al. 2020; Dobson et al. 2020; Gibb et al. 2020a; Carlson et al. 2021; Plowright et al. 2021; Bates et al. 2021; Glidden et al. 2021).

One such study was recently published by Johnson *et al*. (2020), who found that mammal species classified as endangered by the International Union for Conservation of Nature (IUCN) shared relatively fewer viruses with humans (i.e., *zoonotic viruses*, or *zoonoses*). In addition, they found that species that are more globally abundant had significantly greater total zoonotic virus richness than less abundant mammals. These findings connect into a much broader body of literature about the uneven distribution and drivers of viral diversity in animals (Nunn et al. 2003; Lindenfors et al. 2007; Rifkin et al. 2012; Luis et al. 2013; Olival et al. 2017; Albery et al. 2021b, 2021a), and are underscored by similar results, such as an older study on primates that found a similar correlation between threatened status and decreased parasite diversity (Altizer et al. 2007) and a more recent global study that found human disturbance preferentially increases the abundance of species that act as zoonotic reservoirs (Gibb et al. 2020b).

Together, this body of research points to interesting questions about *why* endangered species might be hosts of fewer zoonotic viruses. The most popular explanation so far is the *exposure hypothesis*: any given species might have roughly the same pool of potential zoonotic pathogens, but perturbation and anthropogenic stressors might lead to higher intraspecific contact rates and transmission, higher viral shedding, and more human-wildlife contact. Plausibly, this might actually lead to a *higher* proportional rate of zoonotic emergence in endangered species, if the same factors driving zoonotic emergence are driving their endangered status. However, most existing work hypothesizes that these species fall on the lower end end of a continuum, where declining species unable to tolerate disturbance are less likely to be zoonotic reservoirs, and invasive, weedy species that thrive alongside humans will be less threatened and share more viruses with people (Johnson et al. 2020; Albery et al. 2021b; Plowright et al. 2021). One recent study focused on this latter subset of species that are adapted for life in urban environments (Albery et al. 2021b), which do indeed have a higher average richness of known zoonotic pathogens. Surprisingly, they found instead that species in urban environments are both better studied and have a higher total pathogen diversity than non-urban species, and that the appearance of higher zoonotic pathogen richness is actually explained by these two factors. This finding points to two alternate explanations that can act in combination: the *bias hypothesis* suggests that patterns of viral diversity (zoonotic or otherwise) are highly reflective of research interest and pathogen discovery effort (Wille et al. 2021), while the *susceptibility hypothesis* suggests that there might be differences in the size of species’ underlying pathogen pool (zoonotic or otherwise) driven by interspecific variation in immunology.

Both of these hypotheses are relevant to patterns of viral diversity in endangered taxa. Bias effects could operate in either direction: endangered species might be undersampled both due to rarity and logistical challenges; conversely, infectious disease might be a topic of disproportionate research interest for these species, especially if many are kept in zoos, captive breeding programs, or sanctuaries where veterinarians regularly monitor for disease. However, there could also be underlying biological mechanisms that drive both extinction risk and susceptibility to pathogens, particularly given the relationship between immune investment, life history, and a number of other collinear traits (e.g., trophic level, population density, and geographic range size) that all correlate with pathogen diversity and zoonotic emergence (Plourde et al. 2017; Albery & Becker 2021). Extinction risk also tightly correlates with these same traits (Purvis et al. 2000) and more broadly with species’ “pace of life” (Zaldívar et al. 2004; Ripple et al. 2017; Hernández-Yáñez et al. 2022). All of these patterns might be compounded by the unique features of specific groups of mammals. For example, primates face a high extinction risk and share many viruses with humans given their close phylogenetic relatedness; similarly, many bats are at high risk of extinction, especially in hotspots of environmental change where their role as unique zoonotic reservoirs has been well-studied (Olival et al. 2017). As these examples highlight, it is difficult to disentangle the relationship of susceptibility, exposure, and bias in driving apparent patterns of both total and zoonotic pathogen diversity.

Here, we attempt to separate these competing mechanisms and explain previous findings that endangered mammals have fewer zoonotic pathogens (Altizer et al. 2007; Johnson et al. 2020). To do so, we combined threat status as defined by the IUCN Red List with an open database of host-virus associations (Carlson et al. 2022) and examined both data availability and estimates of viral diversity in each species relative to their endangered status; in doing so, we expand beyond prior studies that focus only on zoonotic viruses or on specific host taxa to consider the full set of viruses that endangered species are known to host across mammals (Johnson et al 2020; Altizer et al, 2007). Using multiple, simple approaches to test causal relationships, we attempt to isolate (1) the role of research effort in driving both total and zoonotic viral diversity; (2) the covariance between the two; and (3) the underlying biological mechanisms that might connect both viral diversity measures to species’ vulnerability to extinction.

## Methods

### Data

#### Host-virus data

To quantify species-level viral richness, we used the Global Virome in One Network (VIRION) database, the most extensive open resource currently available that catalogs known species interactions between vertebrates and their viruses (Carlson et al. 2022). VIRION combines data from several sources, including a manually reconciled core database called CLOVER (Gibb et al. 2021), data aggregated by the Global Biotic Interactions (GLOBI) database (Poelen et al. 2014), and user-submitted data from viral sequences submitted to the National Center for Biotechnology Information (NCBI)’s GenBank database. In its publication release, the database compiles over 23,000 unique interactions among 9,521 viruses and 3,692 vertebrate host species; these data are also updated automatically with new records from GLOBI and GenBank. These data are most complete for mammals, with one in every four mammal species represented in the database and a minimal level of bias in the geographic coverage of host species (though mammal-virus associations are somewhat biased in favor of Eurasia). Here, we used the version 0.2.1 release of the database, subset down to host and viral taxa that have both been resolved with the taxonomic backbone maintained by NCBI, and further limited our analyses to virus species that have been ratified by the International Committee on Viral Taxonomy.

#### Endangered species data

To classify mammal conservation status, population trends, and data deficiency, we followed the threat levels reported by the 2021 IUCN Red List (IUCN, 2021). The IUCN classifies species into one of eight categories: Data Deficient (DD), Least Concern (LC), Near Threatened (NT), Vulnerable (VU), Endangered (EN), Critically Endangered (CR), Extinct in the Wild (EW), and Extinct (EX). These classifications are made primarily on five criteria: small population size, recent decline in population size, small geographical range area, reduced geographical range area, and probability of near-term extinction. For our analysis, we distinguished “endangered” (Endangered or Critically Endangered) from “non-endangered” (Least Concern, Near Threatened, and Vulnerable). Here and throughout the text, endangered in the lowercase refers to this category, while the specific status itself is capitalized; in figures and tables, endangered always refers to the aggregated category, and two-letter abbreviations are used for the specific IUCN Red List classifications. Mammal species classified as “Extinct” or “Extinct in the Wild” were not included in analyses, as information on the viruses of these species is expected to be inherently limited.

Though most scientific names for each species matched between the IUCN and the VIRION database, there were 103 species listed in the VIRION database that were not found in the IUCN records. VIRION records were attributed to corresponding species’ records in the IUCN data for 55 of these differentially labeled species. The remaining 48 species were left unrecorded, as the IUCN Red List had no official records for those species. Following Johnson *et al*., an additional 14 domestic mammal species were manually assigned “least concern” and “increasing” status.

### Analysis

We used two approaches to investigate the relationship between conservation status and viral diversity. The first, a path analysis, allowed us to explore complex interrelationships among variables while explicitly accounting for covariance structures among those that are structurally non-independent (e.g., data deficient status and endangered status are mutually exclusive). The second approach, a mixed-effects regression, allowed us to both incorporate spatial random effects and to model viral richness without violating normality assumptions, but this method collapses the network of causality into two sequential models (i.e., total viral diversity is used as a predictor in the zoonotic virus diversity model, and all other relationships are treated as independent and the same variables are tested in both models). By running both analyses, we were able to test the sensitivity of our result to different assumptions and whether different confounders might explain the effect of endangered status.

#### Path analysis

As a primary approach, we used path analysis to explore the complicated interrelationships among sampling bias, conservation status, and viral discovery. Using the semPlot package in R, we fit a Levain path analysis with several arguments of explanatory and response variables. We generated three binary variables from the IUCN data: the first (“endangered”) reports whether a species is categorized as either Endangered or Critically Endangered; the second (“Decreasing”) reports declining population trends; the third defines species as “Data Deficient.” Our full path model is given in Figure 2 and is based on a set of assumptions that: (1) endangered species may be deliberately better studied (with research effort here measured as PubMed citation counts for a species’ binomial name, as a proxy for total volume of related scientific literature); (2) research effort reduces the chance a species ends up listed as Data Deficient and thus increases the rate of viral discovery; (3) several variables are structurally inseparable, due to the IUCN process or the fact that zoonotic viruses are a subset of total virus counts; and (4) the causal relationships of interest are between conservation status (endangered status, data deficient status, population trends) and viral diversity (both total and zoonotic) while accounting for points 1 through 3. We restricted our analysis to species with at least one virus recorded in VIRION and used a log_10_(x+1) transform on total viral richness, zoonotic viral richness, and citations.

#### Spatial regression analysis

As a secondary approach, we reproduced a modified version of a regression analysis previously used to test whether urban-adapted mammals are more likely to exchange viruses with humans (Albery et al. 2021b). The regression approach (i.e., generalized linear mixed models) treats the number of (total or zoonotic) viruses as a negative binomial distributed variable and includes a log-transformed effect of total richness as a predictor for zoonotic virus richness. The model again treats endangered species, Data Deficient status, and decreasing population trends as binary predictor variables, and also controls for several additional axes of biologically meaningful variation, including host phylogeny (through order-level effects for any group with at least 20 points), life history (quantified using the first two principal components generated by a previous analysis of life history variation in host species (Plourde et al. 2017)), domestication status, geographic range size, and other latent spatial variation. The last of these effects is estimated as a spatial random effect using the integrated nested Laplace approximation (INLA), applied to the centroid of species’ official IUCN range maps. We again restricted our analysis to species with at least one virus recorded in VIRION, as well as complete predictor data, limiting the analysis to 888 total species; citation counts and total viral richness were ln(x+1) transformed. For more details on this approach, see Albery *et al*. (2021b).

## Results

### Overall patterns

Out of the 5,876 Red Listed species in our dataset, only 1,276 species (22%) had any recorded viruses in VIRION. Data availability had a striking correspondence to conservation status, with the proportion with any known viruses decreasing steadily from species listed as Least Concern (28%) to Critically Endangered (14%); only two Extinct in the Wild species had any known viruses (Père David’s deer, *Elaphurus davidianus*; and the scimitar oryx, *Oryx dammah*), as did only one fully Extinct species (the aurochs, *Bos primigenius*). Moreover, nearly all species (98%) listed as Data Deficient have no known viruses (Table 1). Of the species present with any viral associations recorded in VIRION, species with a higher level of threatened status had progressively lower viral richness and zoonotic viral richness, as did species listed as Data Deficient, a fact that is particularly apparent in the long tail of the data (Figure 1).

**Table 1.** The distribution of viral association data available for species, by Red List status. The proportion of zoonotic viruses is relatively constant across groups, and variation between them in the availability of zoonotic richness data is mostly driven by the overall effect of sampling.

| **IUCN** | **LC** | **NT** | **VU** | **EN** | **CR** | **DD** |
| --- | --- | --- | --- | --- | --- | --- |
| Total number of species | 3,338 | 370 | 556 | 542 | 225 | 845 |
| Number of species with any viruses recorded in VIRION | 921 | 87 | 116 | 99 | 32 | 18 |
| Proportion of species with at least one virus known | 27.6% | 23.5% | 20.9% | 18.3% | 14.2% | 2.1% |
| Number of species with any zoonoses recorded in VIRION | 800 | 70 | 101 | 87 | 30 | 15 |
| Proportion of zoonotic reservoirs (out of all species with any known viruses) | 86.9% | 80.5% | 87.1% | 87.9% | 93.8% | 83.3% |

**Figure 1.**
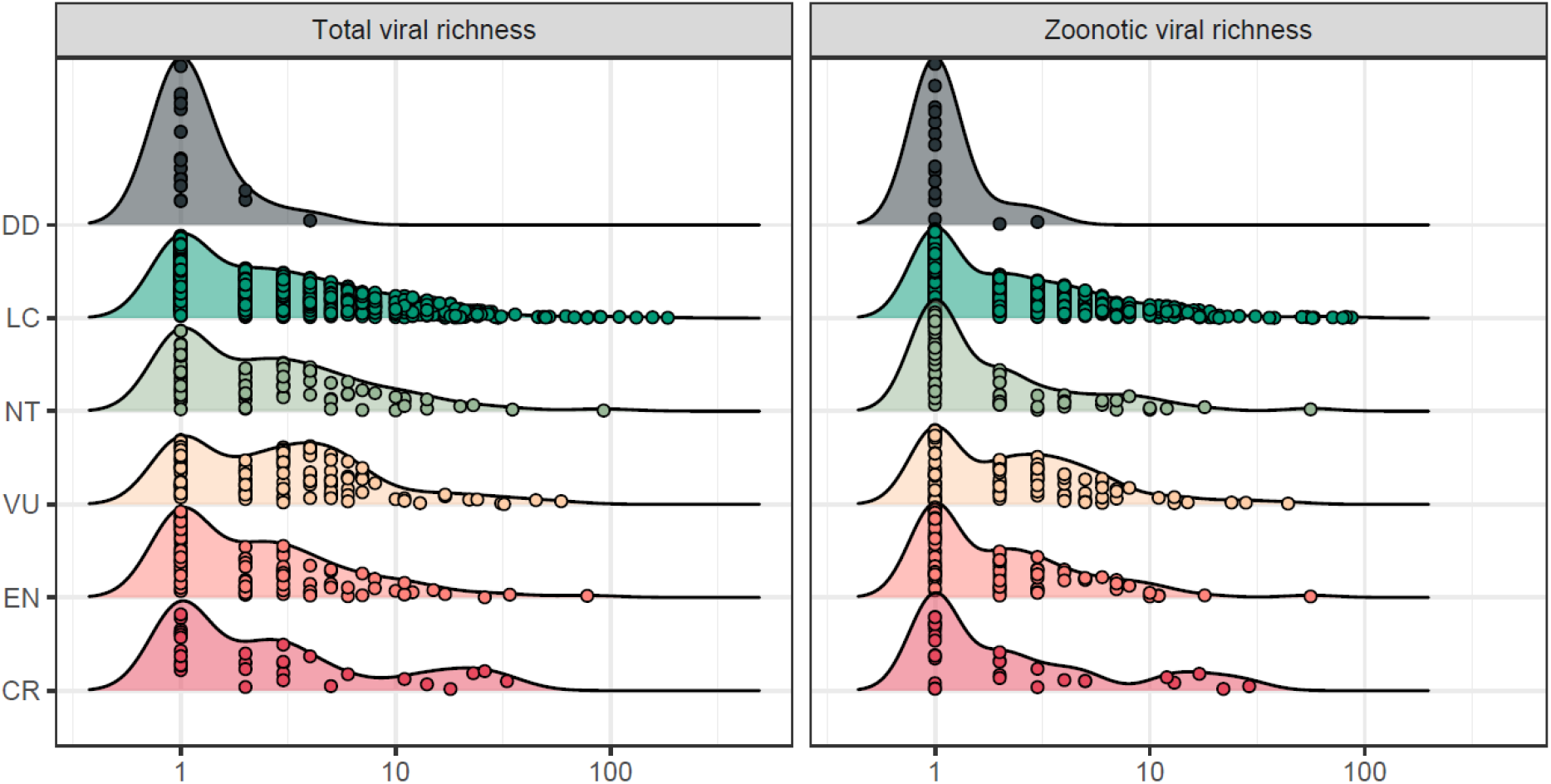
Distribution of virus richness by IUCN status for extant species. (Axes are log-transformed, and zero values are not shown but are described in Table 1; points represent species values and are jittered vertically to show the distribution of raw data.)

### The path model

The path analysis revealed that, even accounting for complex inter-relationships among predictors, endangered species are better studied (i.e., have higher total citation counts) yet have significantly fewer viruses (Figure 2; Table 2). The significant negative effect of endangered status on total viral diversity was the only significant relationship between conservation status and viral diversity; even Data Deficient species did not have fewer viruses than expected accounting for research effort (citations). Unsurprisingly, endangered species were better studied, understudied species were more likely to be listed as Data Deficient, and better-studied species had more total viruses and more zoonotic viruses. Population trends had no significant effect on any variables.

**Table 2.**
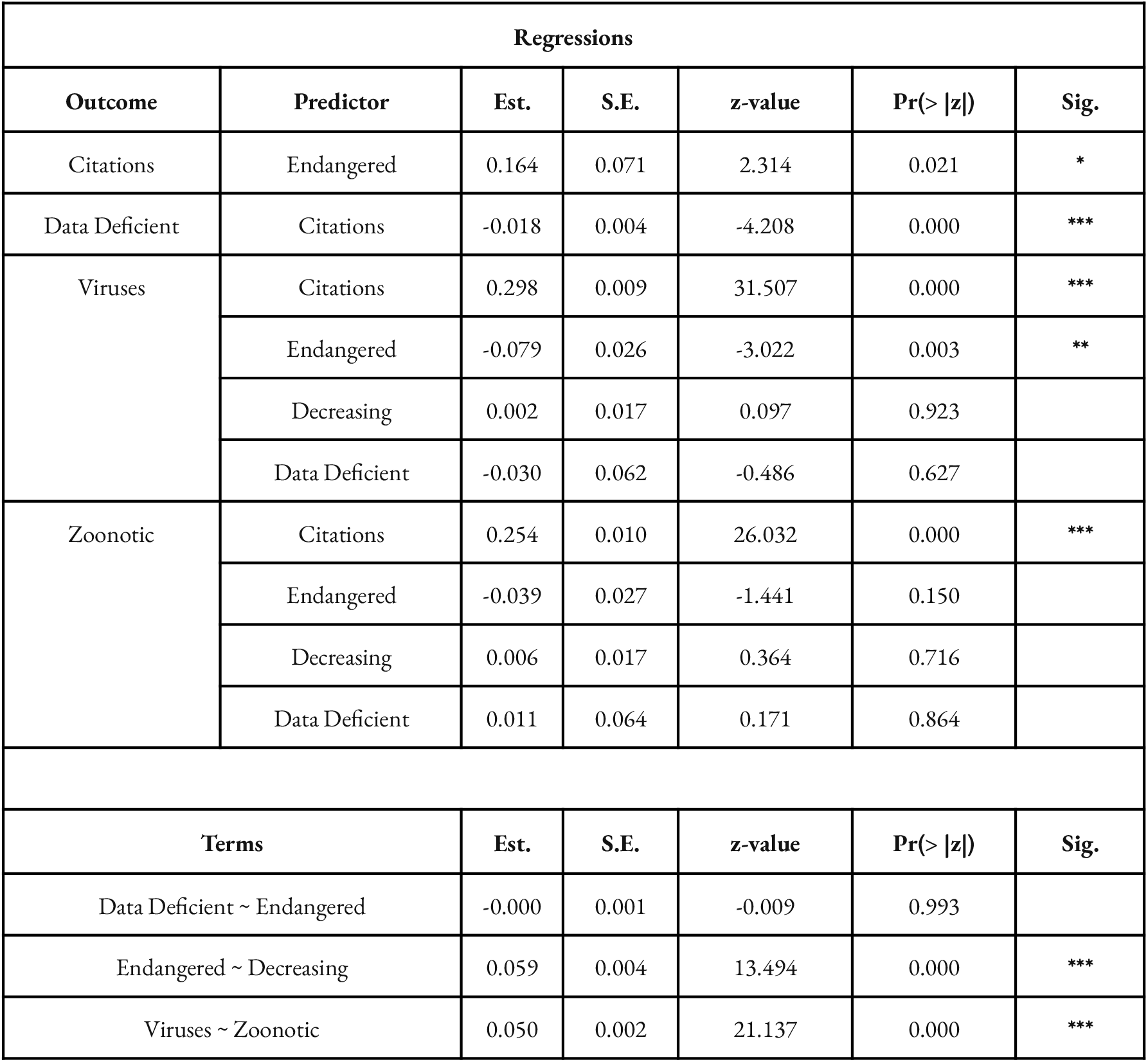
Path analysis coefficients corresponding to relationships in Figure 2. (Significance codes: * p < 0.05; ** p < 0.01; *** p < 0.001)

**Figure 2.**
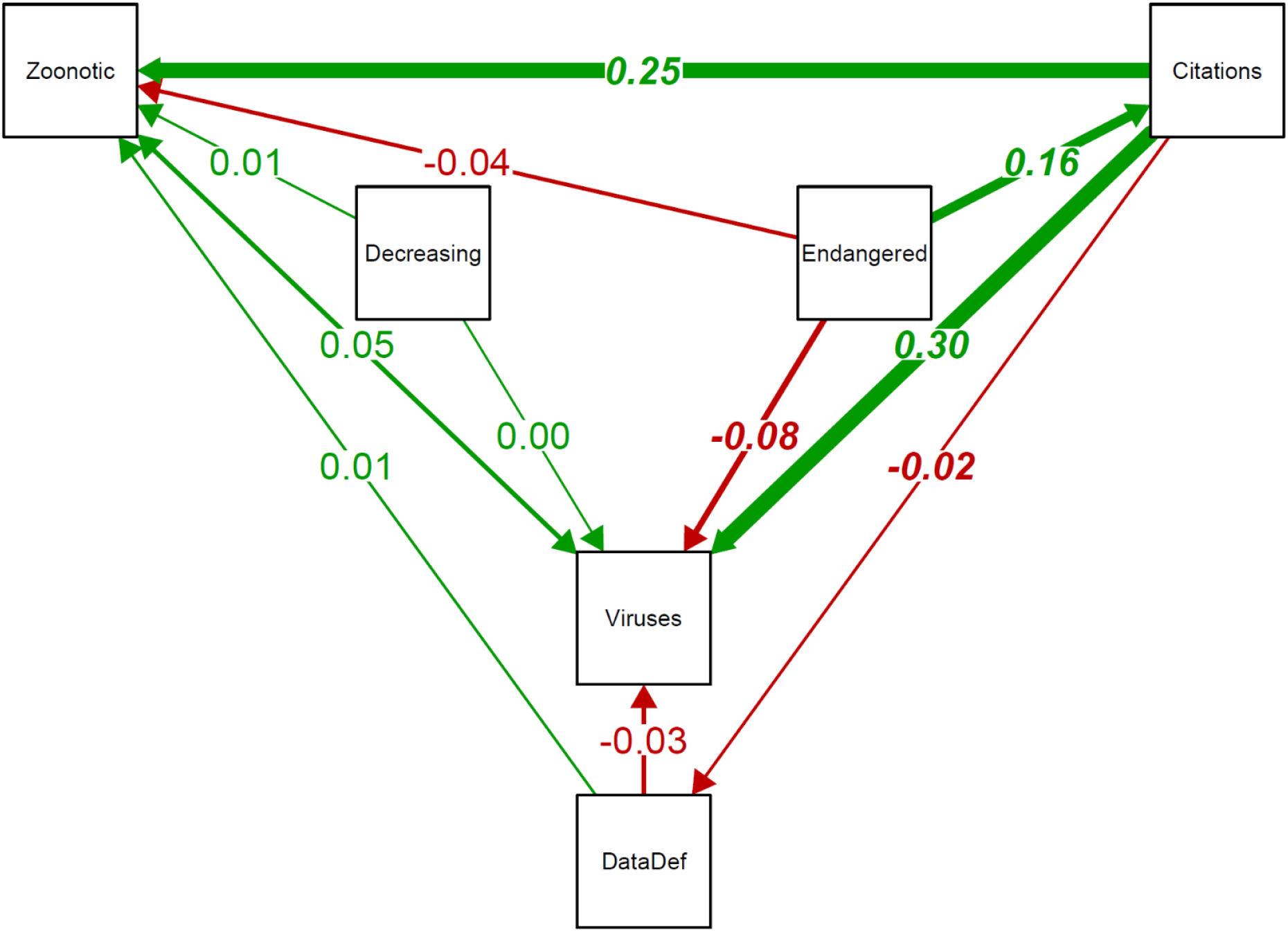
Path analysis revealed that endangered mammals have significantly fewer known viruses than expected (but not fewer known zoonotic viruses) given research effort. Additionally, neither Data Deficient status nor population trends predicted either total or zoonotic viral diversity after accounting for research effort. Arrows denote hypothesized causal relationships with red lines representing negative effects and green lines representing positive effects. Arrow widths are proportional to estimated effects (values mid-line); significant effects are shown in bold italics.

### The regression models

The regression models also found that endangered species had significantly fewer viruses than expected (Figure 3), after accounting for total research effort, spatial patterns of sampling bias, and other interspecific variation like taxonomic and life history differences. The effect of Data Deficient status on species’ viral diversity was strong but insignificant after accounting for the strong and highly-resolved effect of citation counts. Other patterns (e.g., the first axis of life history variation predicted zoonotic richness; primates had a much higher number of zoonotic viruses, and bats had a higher total number of both overall viruses and zoonotic viruses) suggested that the model was detecting other facets of biological variation, but these confounders did not explain the additional effect of endangered status.

**Figure 3.**
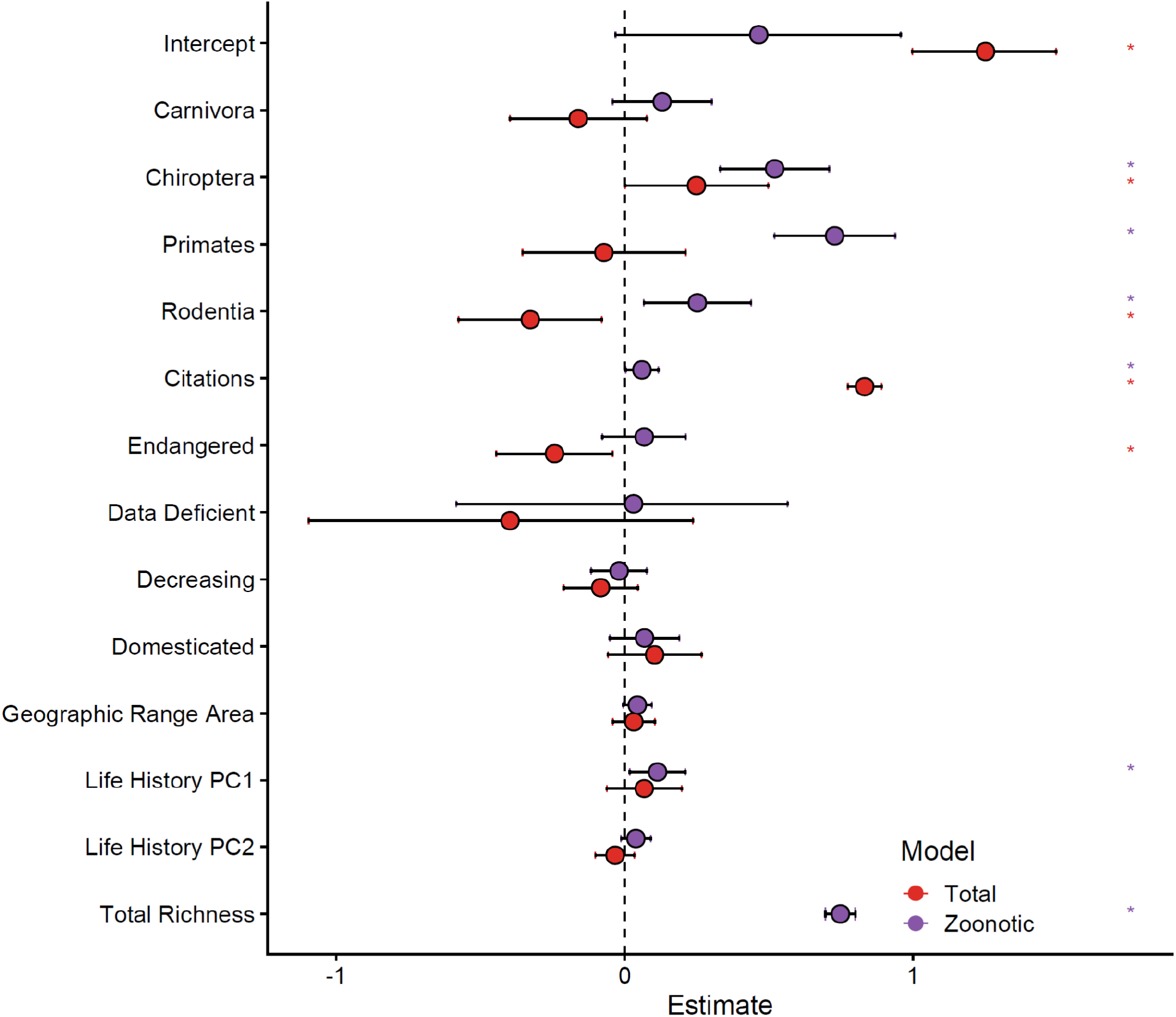
Estimated effects (coefficients and 95% confidence intervals) in the regression model, with total virus richness (red) and zoonotic virus richness (purple) as the outcome variables. Stars indicate significant effects (p < 0.05). Several variables, not listed here, are dropped from the model automatically, including all other mammal order effects and the effect of decreasing population trends. (Artiodactyla is used as the reference group for order-level effects.)

## Discussion

Using multiple statistical approaches, we were able to assign evidence for or against a range of hypotheses about why endangered species have fewer known zoonotic viruses. We found strong evidence in favor of bias effects (i.e., species with Data Deficient status are poorer studied and so have fewer known viruses, including fewer zoonotic viruses), and we found no significant evidence for exposure effects (i.e., the relationship between total and zoonotic virus richness was strong and consistent between endangered and non-endangered species, and increasing population trends had no effect on zoonotic virus diversity). These findings are perhaps unsurprising, given previous work suggesting that patterns in zoonotic virus richness tend to be driven by the total number of viruses known from either specific taxa or broad taxonomic groups (Mollentze & Streicker 2020; Albery et al. 2021b). However, to our surprise, we found a strong relationship between endangered status and total viral diversity that was not explained by either the proxies we used for research effort (i.e., geographic random effects and citation counts, which were actually *higher* on average for endangered species) and underlying host biology that might covary with endangered status (i.e., geographic range size, pace of life, domestication, or taxonomy). Even accounting for all of these factors, endangered species have fewer known viruses.

This pattern might be indicative of other latent biological mechanisms we were unable to test, such as viral coextinctions that take place along the road to host extinction. Although this phenomenon has been studied extensively in macroparasites (Altizer et al. 2007; Farrell et al. 2015, 2021; Carlson et al. 2017; Herrera et al. 2021), this mechanism is generally considered less of a risk to viruses, which are more often host generalists (Harris & Dunn 2013); the full coextinction of a viral species has only rarely, if ever, been documented before (though, see (Das et al. 2020) for a related phenomenon). Our best explanation of this pattern is instead that it reflects an additional layer of complication in the relationship between research interest and viral discovery. Some of this might be driven by passive effects—rare species are less likely to be found when researchers are sampling an entire group (e.g., mist-netting bats)—though the lack of a relationship between population trend and viral diversity runs counter to this idea. More likely, researchers may be less inclined to sample these species for pathogens due to a mix of logistical barriers (e.g., obtaining additional permits or restrictions on sampling in protected areas) and active prioritization (e.g., disease researchers who want to include terminal sampling in their study design, or consider these species challenging to find in the wild, are likely to focus on less threatened species). If true, this is an important dimension of sampling bias that has not previously been considered, with clear ramifications for both conservation and human health: researchers may be less likely to identify pathogens that threaten the survival of these species or that could someday pose a risk to humans.

These ideas point to several important directions for future work. First, researchers might consider reproducing recent analyses of the pace of viral discovery (Wille et al. 2021; Gibb et al. 2022) and testing whether a change to a species’ IUCN Red List status has a downstream effect on research effort, further interrogating our hypothesis that endangered species are subject to an additional and unique form of sampling biases. Further work on the relationship between extinction pressure and disease emergence can also move beyond viral diversity—a crude metric for zoonotic emergence that only minimally captures the transmission process—and examine whether infection prevalence in hosts and shedding into the environment is higher in species facing different kinds of anthropogenic threats (Becker et al. 2021). Finally, we suggest that researchers might consider deliberate efforts to better inventory and study the viral fauna of endangered species, both because of the actionable concerns we raise above, and simply to understand these species better. Despite the logistical challenges of studying these species, as noted above, closer viral surveillance of endangered species should be feasible; for example, one recent study of nearly 100,000 bat samples found no statistically significant effect of terminal sampling on a higher rate of coronavirus detection, and ability to detect virus RNA did not vary strongly across many sample types (Cohen et al. 2022). Collection of non-lethal samples such as oral and rectal swabs, or whole blood or sera, could help further characterize viral assemblages in threatened species—in addition to bolstering voucher collections and providing other conservation-relevant data, like estimates of species’ genetic diversity. In cases where direct sampling still presents logistical or ethical challenges, non-invasive sampling through testing of feces or urine—as sometimes already used for pathogen surveillance in bats and primates (Köndgen et al. 2010; Giles et al. 2021)—represents another avenue to characterize viral communities and their zoonotic potential in these species; recent diagnostic advances also now allow for pathogen risk to be evaluated from environmental DNA (eDNA) or invertebrate-based DNA (iDNA) (Alfano et al. 2021). Lastly, analyses of previously collected voucher specimens and individual samples in museum collections could represent another means to improve research effort of these undertested species (Thompson et al. 2021). Considering how to expand this aspect of global viral surveillance is an important step for conservation biologists aiming to foster a deeper understanding of both wildlife and human health.

## Acknowledgements

This work was supported by NSF BII 2021909 and benefits from an open data and code ecosystem maintained by the Verena Consortium (viralemergence.org), an international collaboration working to predict which viruses could infect humans, which animals host them, and where they could emerge.

## Author Contributions Statement

KN, CJC, and SJR conceived the study; KN, CJC, and GFA produced analyses and visualizations; all authors contributed to the conceptualization and writing.

## Data and Code Availability Statement

All code and data for this project is available on Github at github.com/viralemergence/dangerzone

## Conflict of Interest Statement

The authors declare no conflicts of interests.

## Notes

http://www.github.com/viralemergence/dangerzone

### Competing Interest Statement

The authors have declared no competing interest.

